# The Impact of Leishmaniasis on Mental Health and Psychosocial Well-being: A Systematic Review

**DOI:** 10.1101/637132

**Authors:** Malini Pires, Barry Wright, Paul M. Kaye, Virgínia Conceição, Rachel C. Churchill

## Abstract

**Background:** Leishmaniasis is a neglected tropical parasitic disease endemic in South Asia, East Africa, South America and the Middle East. It is associated with low socioeconomic status (SES) and responsible for considerable mortality and morbidity. Reports suggest that patients with leishmaniasis may have a higher risk of mental illness (MI), psychosocial morbidity (PM) and reduced quality of life (QoL), but this is not well characterised. The aim of this study was to conduct a systematic review to assess the reported impact of leishmaniasis on mental health and psychosocial wellbeing.

**Methods:** A systematic review of the literature was carried out. Pre-specified criteria were applied to identify publications including observational quantitative studies or systematic reviews. Two reviewers screened all of the titles, abstracts and full-studies and a third reviewer was consulted for disagreements. Data was extracted from papers meeting the criteria and quality appraisal of the methods was performed using the Newcastle-Ottowa Scale or the Risk of Bias in Systematic Review tool.

**Results:** A total of 14 studies were identified from 12,517 records. Nine cross-sectional, three case-control, one cohort study and one systematic review were included. Eleven assessed MI outcomes and were measured with tools specifically designed for this; nine measured PM and 12 measured QoL using validated measurement tools. Quality appraisal of the studies showed that six were of good quality. Cutaneous leishmaniasis and post kala-azar dermal leishmaniasis showed evidence of associated MI and PM including depression, anxiety and stigma, while all forms of disease showed decreased QoL. The findings were used to inform a proposed model and conceptual framework to show the possible links between leishmaniasis and mental health outcomes.

**Conclusion:** There is evidence that leishmaniasis has an impact on MI, PM or QoL of patients and their families and this occurs in all the main subtypes of the disease. There are however large gaps in the evidence. Further research is required to understand the full extent of this problem and its mechanistic basis.

**AUTHOR SUMMARY:** Leishmaniasis is a parasitic disease prevalent in many low-and middle-income countries worldwide. In this study the authors looked for evidence as to whether leishmaniasis affected the mental health and quality of life of patients. To conduct the review, a wide search of the literature was conducted, where a total of 14 full articles were included and analysed. It was found that different forms of leishmaniasis (visceral leishmaniasis, cutaneous leishmaniasis and post kala-azar dermal leishmaniasis) do cause a significant impact on patients’ mental health and quality of life through societal factors such as stigma, lack of knowledge, culture and low self-esteem among others. However, no evidence of biological mechanisms was found linking leishmaniasis to mental illness or decreased quality of life. Despite being a very incapacitating disease physically, leishmaniasis also leads to mental illness and decreased quality of life, and should therefore be a priority on the global health agenda for both researchers and policy makers.

## Introduction

Leishmaniasis is a neglected tropical disease (NTD) caused by multiple species of *Leishmania* parasites and transmitted by the bite of female sand flies. It is endemic in 98 countries, and is mostly concentrated in low-and middle-income countries in South Asia, East Africa, South America and in the Middle East (1). The disease presents in different forms depending on the species and geographical location. Visceral leishmaniasis (VL; also known as kala-azar) presents with fever, weight loss, hepatosplenomegaly and may have neurological manifestations (2). If untreated, it has a fatality rate of over 95% (3). Post kala-azar dermal leishmaniasis (PKDL), occurring as a consequence of VL can cause erythematous or hypopigmented macules, papules, nodules and patches (4). Cutaneous leishmaniasis (CL) patients present with plaques, nodules and / or ulcers and, in the case of mucocutaneous leishmaniasis (MCL), symptoms manifest on the mucous membranes of the nasal and oral cavities and surrounding tissues (5). These forms of leishmaniasis invariably leave visible disfiguring lesions and lifelong scars on the skin (6,7).

Although not fatal, CL lesions have been described in the literature as a source of distress and discomfort. Such visible manifestations have been linked to stigmatizing attitudes that could potentially lead to social stigmatisation, (6,9) and psychosocial morbidity (PM). For example, in Afghanistan, the incorrect belief that the disease can be transmitted by human contact results in social exclusion that can range from not sharing utensils to severe physical and emotional isolation (10). In some cultures, women are considered unfit for marriage or are separated from their children when they have the disease(10,11). A study involving high school students in Morocco showed awareness of the stigma attached to CL with self-stigmatization including feelings of shame, embarrassment, depression, and self-contempt (6).

Although inflammation of the liver and spleen are the most well-documented clinical manifestations of VL, inflammation of the nervous system has also been reported (Lima et al., 2003, Marangoni et al., 2011). Neurological manifestations in human VL include peripheral and cranial nerve dysfunction, neurological tremors, meningitis, seizures, paresis, and haemorrhagic stroke (12–18). In addition, naturally infected dogs with canine VL as well as rodent models of VL show neuroinflammation (19,20). Despite these findings and research showing close links in general between neuroinflammation and mental health problems (21), there is limited evidence of a direct link between the neuroinflammation present in leishmaniasis and mental health problems.

There is a growing body of evidence suggesting a causal link between systemic and localised inflammation and depression in other mental health disorders (Stewart et al., 2009; Taraz et al., 2012; Oliveira Miranda et al., 2014; Ford and Erlinger, 2004; Vogelzangs et al., 2016;Furtado and Katzman, 2015) including 2 meta-analyses that show statistically significant raised levels of inflammatory markers in depressed subjects (28,29). Patients with depression often show cardinal features of inflammation, including increased expression of pro-inflammatory cytokines and their receptors as well as an upregulation of acute-phase reactants, chemokines and soluble adhesion molecules in peripheral blood and cerebrospinal fluid (CSF) (25,30,31). In VL, a link between mental illness (MI) and systemic or neuroinflammation been postulated (20).

At the time this review was conducted, there had been no high quality comprehensive systematic review published on possible aetiological associations and the impact of leishmaniasis on mental health and psychosocial wellbeing. A search of the Cochrane Database of Systematic Reviews and the Database of Abstracts of Reviews of Effects (DARE) identified no systematic reviews of the effects of healthcare interventions for the potential mental health consequences of leishmaniasis. A detailed search of PROSPERO indicated that no other systematic reviews were in progress. Subsequently, one scoping review has recently been published (29) making a ‘preliminary assessment of the extent of the literature’ focusing solely on localised cutaneous leishmaniasis and with no quality appraisal for included studies.

The objective of this study was, therefore, to conduct a comprehensive systematic review (SR) of MI, PM and quality of life (QoL) related to all forms of leishmaniasis. Secondary research questions included: i) is there evidence that inflammation resulting from VL is directly associated with MI?; ii) is there evidence for social stigma against people with VL? If so, is this associated with MI?; iii) do cognitive and physical impairments resulting from VL result in MI or PM?; iv) are stigma and discrimination of patients with post kala-azar dermal leishmaniasis and cutaneous leishmaniasis associated with MI?; and v) do co-morbidities in people with leishmaniasis have an association with decreased QoL, increased MI or PM?

From this research, we set out to develop a conceptual model for the association between leishmaniasis, mental health outcomes and PM. This model may serve to prompt further research and debate and to inform health workers, government bodies and the scientific community about the nature of and the mental health implications of leishmaniasis and the unanswered questions surrounding these associations.

## Methods

The review methodology including the search strategy for this review was published in PROSPERO (32) following CRD (33) and PRISMA (34) guidelines. The recommendations in the Cochrane Handbook (Higgins and Green et al 2011) were adhered to for reporting the review.

### Databases and Search Strategy

The following primary electronic databases were searched after expert advice from experienced information specialists working in this field: MEDLINE, EMBASE, PsycINFO, LILACS, POPLINE, Global Health, IndMED, ArabPsyNet and AfricanIndexMedicus (AIM). There were no restrictions on the date and language of publication. MEDLINE and EMBASE were chosen as comprehensive databases for life sciences and biomedical research. PsycINFO is a robust database which contains psychology-related articles. POPLINE and Global Health are population and public health-related databases. LILACS, IndMED, ArabPsyNet and AIM were searched as they contain literature from geographical locations where leishmaniasis is endemic (32).

The search strategy (S3) was designed to be more sensitive than specific (search strategy included all possible terms to answer the broad PICOS as opposed to one with less descriptors) as potentially unknown exposures could exist, and because of the lack of a previous robust synthesis of this topic. Backward and forward citation tracking was performed for included studies to find any relevant studies not included in the databases. After retrieving the results for each database, citations and abstracts were exported to EndNote to remove duplicates and for screening.

### Rationale for the chosen outcomes

The outcomes were a broad scope of the psychosocial consequences that can result from leishmaniasis. They were based on three concepts: 1) the biological link between the neuroinflammation caused in VL and MI; 2) the social impact of stigma, isolation and discrimination around CL on QoL; 3) any further burden on the patient in terms of QoL or mental health (including impact on their relatives) such as, for example consequences secondary to outcomes related to mental health (e.g. financial or physical co-morbidities).

### Rationale for chosen study designs

Cohort studies, case-controls and cross-sectional studies that quantitatively measured psychosocial outcomes such as mental illness or QoL using a validated tool in this population were included. Systematic reviews (including and in addition to the review by Bennis et al 2018) were also included as a mechanism for identifying relevant publication for screening.

### Study eligibility criteria

The inclusion and exclusion criteria used to select studies is shown in the protocol (32), which used an adapted PICOS framework for observational studies. As this study was examining observational studies and not randomised control studies, there was no “intervention” being studied, and instead, the term “exposure” was used. There was also no comparator in observational studies, so this domain was not used.

### Rationale for chosen population

Due to the broad exploratory nature of this systematic review, all patients with any form of leishmaniasis regardless of age and gender were included. We also included assessments of how the disease has wider impacts on family and community members due, for example, to social stigma and financial strain (35,36).

### Rationale for chosen exposures

The exposures included the social determinants of health associated with leishmaniasis including: poverty, low social economic status, co-morbid infections, social and cultural norms, as well as the physical and psychosocial factors that could lead to a decrease in QoL, PM and MI.

### Data extraction

Titles and abstracts were downloaded onto EndNote and de-duplicated. These were then transferred onto an excel spreadsheet and screened independently by two reviewers (MP and VM). Any disagreement between these two authors were resolved by a third reviewer (RC). Screening codes for titles and abstracts were also created. After selecting the included studies, a standard form as used to extract the relevant data. Extracted information included: study setting; study design; year; time period; study population; sample size; inclusion rate and attrition (where relevant e.g. for cohort studies); age range of participants; mean age; sex ratio (M/F); outcome (e.g. anxiety or depression); statistical test; measurement tool and main findings.

### Data Synthesis and Quality Assessment

A narrative synthesis of the findings from the included studies was performed. This was organised according to characteristics of the studied population; outcome and how these were measured as well as measures of effect. It was expected that the studies would not be similar enough to conduct a meta-analysis due to the heterogeneity of the primary studies included in the review e.g. cutaneous, visceral, PKDL; different countries, different studied populations, different outcome measures and different study designs).

### Quality Assessment and Risk of Bias

The tool to assess quality and risk of bias for the cohort and case-control studies was the Newcastle-Ottawa Scale (NOS) (37) as suggested in the Cochrane Handbook (38). For cross-sectional studies, an adapted version of the NOS was used (39). For any SRs identified during the selection process, the ROBIS tool for assessing the risk of bias was used. The tool is comprised of three phases: 1) assess relevance, 2) identify concerns with the review process, and 3) judge risk of bias (40).

### Methodological Quality Appraisal of Studies

The methodological quality of the studies was assessed for the 13 primary studies using the NOS risk of bias tool, as described above; the systematic review (Bennis et al., 2018)was assessed using ROBIS. An adapted version of NOS was used to appraise the quality of the cross-sectional studies (39). The case-control studies and cohort study were appraised using the original version of NOS.

### ROBIS

ROBIS is a very comprehensive tool used to assess the risk of bias in systematic reviews, and not in primary studies (40). It was used to assess the risk of bias in the systematic review that met our inclusion criteria (41).

## RESULTS

### Study Characteristics

A total of 14 publications (consisting of 13 full articles and 1 conference proceeding) met all of the inclusion criteria for this systematic review (41,42,51–54,43–50) (S2). All were independent studies, except two (42,43) where one was the baseline study, and the other the follow-up study, and published between 2004 and 2018. Three studied VL (42,43,46); 10 studied CL (41,44,45,48,49,51–55) and one PKDL (50) (Table 1). All studies were performed in LMICs (Table 1).

**Table 1.** General Characteristics of the Included Studies.

Summary of data obtained from data extraction process.
| **Author**<br>**Year**<br>**Country** | **Study**<br>**Design** | **Sample (population)** | **Leishmani**<br>**asis**<br>**Subtype** | **Mental health**<br>**consequence**<br>**measured**<br>**(outcome)** | **Sample**<br>**size** | **Age in years**<br>**(mean±SD**<br>**or range (if**<br>**mean n/a)** | **Gender ratio**<br>**(Female %)** |
| --- | --- | --- | --- | --- | --- | --- | --- |
| Alemayehu et al.<br>2017<br>Ethiopia | Cross-sectional | HIV infected patients receiving antiretroviral therapy (ART)- with and without VL coinfection at four different sites in Northwest Ethiopia | HIV-VL | QoL with WHOQoL-HIV instrument | 590 | 34.5 (± 7.4)-<br>HIV-VL patients<br>36.4 (±8.8)-<br>HIV-only<br>patients | 3.2%-<br>HIV-VL patients<br>61.7%-<br>HIV-only patients |
| Alemayehu et al.<br>2018<br>Ethiopia | Prospective cohort | Same as above (6-month follow-up) | HIV-VL | QoL with WHOQoL-HIV instrument | 581 | 34.5 (±7.7)-<br>HIV-VL patients<br>36.4 (±8.9)-<br>HIV-only<br>patients | 5.3%-<br>HIV-VL patients<br>61.7%-<br>HIV-only patients |
| Bennis et al.<br>2018<br>Several Countries | Systematic Review | Patients or their relatives experiencing a skin condition linked to LCL in any country. | LCL | PM, QoL and MI. No quality appraisal tool used | n/a* | * | * |
| Chahed et al.<br>2016<br>Tunisia | Cross-sectional | CL patients with scars selected randomly in primary health centres | CL | QoL, PM and MI with IPQ-R, PSLI and WHOQOL-26 | 41 | 12-53 | 100% |
| de Castro Toledo et al.<br>2013<br>Brazil | Cross-sectional | Patients with confirmed diagnosis of CL with a minimum of 4 years of school education | CL | QoL with DLQI | 20 | 45.6 | 15% |
| Govil et al.<br>2018<br>India | Cross-sectional | People living in villages where VL is endemic. | VL | PM with KAP structured questionnaire | 353 | 41.7 ($\pm 17.1$) | 44.5% |
| Handjani and Kalafi 2013<br>Iran | Cross-sectional | Family members | CL | QoL with FDLQI | 5 | 42 | 27% |
| Honório et al.<br>2016<br>Brazil | Cross-sectional | CL patients at University Hospital of Brasilia | CL | QoL, PM, and MI with WHOQOL-BREF | 44 | 51.8 ($\pm 11.6$) | 54.5% |
| Layegh et al.<br>2017<br>Iran | Cross-sectional | Children with CL with in dermatology clinics. | CL | QoL and MI with CDLQI, CDI and STAIC | 42 | ** | 69% |
| Pal et al.<br>2017<br>India | Case-control | Outpatient and inpatient population. | PKDL | QoL with DLQI | 188 | 27.4- patients<br>29.5 healthy controls | 45.6%- patients<br>44.8%- controls |
| Simsek et al.<br>2008<br>Turkey | Cross-sectional | Women selected randomly from cluster sampling from households. | CL | MI with SCID-I | 270 | 33.3+9.4 | 100% |
| Turan et al.<br>2015<br>Turkey | Case-control | Children and adolescents with CL and healthy controls, and their respective mothers. | CL | MI and QoL with CDI PedQL-C, PedQL-P, STAIC | 54 | 12.0 ($\pm 3.2$) | 46.3% |
| Vares et al.<br>2013<br>Iran | Cross-sectional | patients >16 years-old with CL | CL | QoL and psychosocial morbidity with DLQI | 124 | 36.9 ($\pm 14.9$) | 62.9% |
| Yanik et al.<br>2004<br>Turkey | Case-control | Patients with Active CL and healthy controls from a leishmaniasis treatment centre. | CL | QoL, MI and PM with HAD, BIS and DQL | 99 | 18.8 ($\pm 5.9$) | 50.5% |

Not applicable \*
Not mentioned \*\*

### Population and Demographics

Three studies were on VL (42,43,46); 10 studied CL (41,44,45,48,49,51–55); and Pal et al., 2017 studied PKDL (50). Of the VL studies, two were about HIV patients co-infected with VL (42,43), which was part of the inclusion criteria for the population.

The populations measured in the studies vary greatly in age. Ages of the participants in the studies ranged from the ages of 7 to 80 years, according to the values reported in the studies. Two of the studies were in children only (49,52). Most of the studies had both female and male participants although two (44,51) had an only female population. The combined number of study participants in the 14 studies was 2565. Sample sizes of the primary studies ranged from 20 (45) to 620 (42).

Female to Male ratio in percentage of female was 57% showing that more women were studied over-all (excluding the review) (41).

All studies, including the systematic review were performed in low-and middle-income countries, according to the World Bank’s listing of these countries; Ethiopia is classified as low-income, India as a lower-middle income; and Brazil, Turkey and Iran as upper middle-income (56).

### Study Design

Nine studies were cross-sectional (42,44–46,48,49,51,53,55), three were case-control (Pal et al., 2017; Turan et al., 2015; Yanik et al., 2004), one was a prospective cohort study (43) and one a systematic review (41). The characteristics of the 14 included studied that met the eligibility criteria were assessed for quantitative synthesis.

### Diagnostic Criteria

The different instruments to measure the outcomes for each of the 14 publications are shown in Table 1.

### Outcomes

#### Visceral leishmaniasis

Two studies by the same authors conducted in Northwest Ethiopia reported the results of a prospective longitudinal study at baseline (42) and followed up six months later (43), measuring QoL in HIV patients with VL (HIV-VL) and HIV patients. At baseline, HIV-VL patients had lower mean scores in all domains of the QoL questionnaire showing poor quality of life. Importantly depression was strongly and consistently associated with all the QoL domains in HIV-VL patients (as it was in the HIV group. The mean (SD) depressive-symptoms scale scores were higher 2.67 (±0.7) for HIV-VL patients compared to HIV patients 1.61 (±0.5) (p = 0.001) (42). After 6 months of treatment with both antiretroviral treatment (ART) and anti-leishmanial drugs, the follow-up study (43), showed there was improvement in all the QoL domains analysed at baseline in both groups. Mean scores for social relationship among co-infected patients were significantly lower compared to the HIV group (p=0.001).

Another study looked at knowledge attitudes and practices (KAP) about VL among adults in a community in India. It was found that 7.6% of the participants agreed with the statement that the incidence of VL in the family should not be disclosed. Forty-three percent reported that the illness affects mental health, causes stress (27%), irritation (3.7%), depression/fear of death (5.8%) and other (16,9%). Almost 74% thought that VL in the family has financial consequences, causes impoverishment. (39.5%) leading to the need for loans (17.6%), forced to sell property (0.7%) and other (15.5%) (46).

#### Cutaneous leishmaniasis

The cross-sectional study by Chahed et al, 2016, measured the QoL in girls and women with CL scars and explored the psychological and psychosocial consequences of CL using the Revised Illness Perception Questionnaire (IPQ-R), World Health Organization Quality of Life-26 (WHOQOL-26) and the Psoriasis Life Stress Inventory (PLSI) in 41 girls and women with CL scars in the Sidi Bouzid region, Tunisia (44). The correlation analyses performed on inter and intra-subscales showed that the emotional representations associated with CL were correlated with a loss of self-esteem feelings of inferiority (r=0.77, p<0.05). The more patients knew about CL, the more pessimistic they got about the prospects of recovery. Patients who had a more coherent perception of CL had stronger emotional reactions (p<0.001). Moderate correlations were found between the total number of body scars and experiences of rejection (r=0.31, p<0.05). The number of body scars had a strong link to experience with stigma. Experiences of rejection and avoidance of stress negatively correlated with age (r=-0.33, p<0.05, and r=-0.31, p<0.05), suggesting that younger women experience social stigma more than older women. The WHOQOL-26 and the PSLI questionnaire results showed that the domains of Social QoL and anticipation avoidance of stress, and social QoL and total stress correlated significantly (r=-0.36, p<0.05 and r=-0.32, p<0.05). The Mental QoL domains, however, did not correlate significantly with any of the PSLI domains.

QoL was measured using the Dermatology Life Quality Index (DLQI) in a cross-sectional study of 20 patients with CL (45). In 70% (n=14), CL resulted in a moderate to large effect on QoL in the work and school domain, and the “symptoms and feelings” domain followed. The domain with the least impact was “personal relationships”.

Handjani and Kalafi, 2013 measured the impact of dermatological diseases on the quality of life of healthy families of patients with skin diseases which included five patients with CL using the 10-item validated Persian version of the Family Dermatology Life Quality Index (FDLQI) questionnaire (55). The FDLQI scores for each of the groups showed that there was no statistically significant difference in QoL of families found between the groups of different skin diseases studied (vitiligo, psoriasis, pemphigus and leishmaniasis). However, due to there only being 5 CL families in this study, there could be lack of statistical power.

The WHOQOL-BREF was used to study QoL, PM and MI of CL patients (48). The psychological and environment domains had the lowest median scores. Forty (90.9%) interviewees presented negative feelings (blue mood, anxiety, despair, depression). Of these, eight (18.18%) reported experiencing such feelings always, 19 (43.18%) very often, nine (20.45%) quite often, and four (9.09%) rarely. 50% were dissatisfied with the support received from family and friends, and in their intimate lives.

Layegh et al., 2017, measured the QoL, depression, and anxiety (MI) in children with CL using the Children’s Dermatology Life Quality Index (CDLQI), Children’s Depression Inventory (CDI), and State-Trait Anxiety Inventory for Children (STAIC) questionnaires (49). This study enrolled 42 children by convenience sampling. The prevalence of low quality of life, state anxiety, and trait anxiety was 57.1, 76.9, and 15.8%, respectively; 32% of patients had depression. Cases of low quality of life (54.1%), state anxiety (56.6%), and trait anxiety (53.8%) were more common with the acute form of leishmaniasis. Low quality of life (70.83%), state anxiety (76.66%), trait anxiety (83.3%), and depression (84.6%) were more prevalent in females. The face was the most common location of involvement in patients with low quality of life (63.3%), state anxiety (70.4%), trait anxiety (83.3%), and depression (54.5%). However, the authors report that no significant difference was found between psychological factors in patients and sex (p > 0.05), acute or chronic type of disease (p > 0.05), presence of any other skin or systemic diseases (p > 0.05), location of lesions (p > 0.05), number of lesions (p > 0.05), and duration of involvement (p > 0.05). No information is available about the statistical tests used. This communication was in the form of a conference abstract (49).

Simsek et al., 2008 conducted a cross-sectional study to assess mental health disorders in women in the region using the Structured Clinical Interview for DSM-IV Axis I Disorders (SCID-I).0.5) (51). Fifteen out of 270 women had cutaneous leishmaniasis (6.1%) of whom 8 (53.3%) had a mental disorder. Women with CL have a 2.13 higher OR (95% CI: 1.25-7.31) compared to women without CL of suffering from a mental disorder mainly depression and anxiety.

Turan et al., 2015 assessed the psychiatric morbidity and QoL in children and adolescents with CL and their parents using the Child Depression Inventory (CDI), the State-Trait Anxiety Inventories for Children (STAIC) and the Pediatric Quality of Life Inventory Parent and Child Versions (PedQL-P and C, respectively) (52). Fifty-four subjects, of mean age 12.0 (±3.2) years of whom 46.3% were female, were matched with 40 healthy controls of mean age 11.5 (±2.3) years, 50% of whom were female. In addition, the mother or other caregiver was included for each child. From 2011 to 2013, subjects were recruited at the paediatrics department of a hospital in Sanliurfa, Turkey and the controls were children receiving vaccinations or undergoing routine health checks, matched for age, gender, and parents’ level of education. Scores for depression were higher in patients compared to the controls and QoL was lower in patients and their mothers. All results were statistically significant (p<0.001 for CDI and p<0.05 for PedQL-P and PedQL-C). However no statistically significant difference in scores for anxiety (STAIC) was found.

Yanik et al., 2004 looked at the psychological impact of CL using the Hospital Anxiety Depression Scale (HAD), the Body Image Satisfaction Scale (BIS) and the Dermatology Quality of Life Scale (DQL) to measure anxiety, psychosocial morbidity and quality of life, respectively. Ninety-nine subjects in 3 equal groups, those with active lesions (n=33), patients with healed lesions (n=33) and a healthy group (n=33) were studied. Results showed that lesions on the face and hands, disease active for over a year, permanent scars and social stigmatisation led to anxiety and depression. Body image satisfaction and quality of life were also decreased. Higher scores were obtained in patients with active CL and scores were also statistically significant in patients with healed scars compared to controls in HAD and BIS. Patients with active lesions had lower QoL scores compared to those with healed scars. The correlations between the subscale of HAD and DQL showed a moderate correlation (anxiety r=0.490, p< 0.001; depression r=0.291, p=0.040). The comparison between the HAD scale and the BIS scale had a significant correlation (anxiety r=0.201, p=0.047; depression r=0.256, p=0.011).

The quality of life in patients with CL using the Dermatology Quality of Life (DLQI) Index was carried out (53). QoL was significantly affected. Highest scores were seen in the symptoms and feelings domains; the lowest effect was observed in the treatment domain of the DLQI. The appearance of the lesion and type of the lesion significantly affected the QoL (p<0.05) as patients with ulcerated lesions had lower quality of life compared to those with nodular (P = 0.003) and plaque lesions (P = 0.005). The activity of the disease, location of the lesions and gender did not affect the scores significantly.

A scoping review on the impact of localised cutaneous leishmaniasis on psychosocial wellbeing has recently been conducted. Eight quantitative studies (44,51–55,57,58) five qualitative studies (6,9,11,59,60) and two mixed-methods (10,61) studies were included in their review. It combines the results of the quantitative studies through narrative synthesis looking at anxiety and depression, low QoL, stigma and fear of scars. The three last cited quantitative studies were picked up by us during the title screen but did not fulfil the abstract or full-article screen criteria for this review, and were thus excluded (refer to protocol for inclusion and exclusion criteria) (32).

Our set of inclusion criteria was different from that of the scoping review carried out by Bennis and colleagues (date), hence only six studies included in their study overlap with this systematic review (44,51–55). The quantitative study by Abazid et al., 2012 showed no mental health or psychosocial outcomes (57); An RCT was found that falls under our exclusion criteria (58). Qualitative studies were not included in our study.

Bennis et al., failed to pick up three studies on CL that were identified and included in this systematic review (45,48,49).

#### Post kala azar dermal leishmaniasis

Pal et al., 2017 assessed the perceptions, stigma and quality of life in patients with PKDL patients compared to healthy controls using the Dermatology Life Quality Index (DLQI) and the 36-Item Short Form Health Survey (SF-36) (50). The type and severity of the lesions were also noted. A range of independent variables were compared. DLQI scores a significant effect on QoL in patients of PKDL with highest impairment found in patients under the age of 20 years (p =0.03) and with type of lesion, where patients with the more severe nodular lesions had poorer scores (p = 0.001). Initiation of treatment for PKDL improved the scores (p = 0.04), while gender, duration and location of the lesions had no impact. The SF-36 showed that mental health, social functioning, body pain and general health were significantly different (p<0.05) in the patients compared to the control group.

#### Risk of Bias assessment

Based on NOS, 6 of the 14 studies were of good quality, 3 were of fair quality and the remaining 4 were of poor quality (S4 Table). The fact that some studies were carried out in LMICs of lower economic status than others did not affect the quality outcome. Alemayehu et al 2017 and 2018 both showed the highest scores among all 14 articles, despite having been carried out in Ethiopia (42,43). Most studies with poor or fair scores were found to have inadequate sample sizes (too small) with no justification (44–49,53). In contrast, recruitment methods for subjects seemed to be justified for all studies. Out of the nine cross-sectional studies, only five (42,45,46,49,53) controlled for at least one confounder. Thus, the remaining four could be severely biased due to confounders such as the age or gender of the participants. Moreover, what contributed further to risk of bias was that five of the studies (45,47–49,53) did not report confidence intervals, which is considered inadequate statistical reporting. However, some important inconsistencies between the NOS and the adapted-NOS, as well as the different study designs, could potentially have had an impact on the overall score and quality rating. The maximum rating that can be given in the adapted NOS-version is 10 stars whereas for the original NOS versions (cohort and case-control studies) Quality rating (S4 Table) does not take into account the overall star rating, but rather the rating per section: “selection”, “comparability” and “exposure”. The comparability section was always assessed in terms of whether the study controlled for specific confounding variables. Gender and age were considered the most important variables to control for.

The ROBIS (S5-S8 Tables) assessment showed that in the review by Bennis et al 2018 the risk of bias is unclear overall. As shown in the assessment, the lack of a previously published protocol made it very difficult to assess the reliability of the review.

## DISCUSSION

The primary research question of this review was: *What is the impact of leishmaniasis on mental health and psychosocial wellbeing?* This systematic review shows preliminary evidence of associations between leishmaniasis and mental health, but also shows several lacunae in the evidence. The systematic review only yielded studies about VL, PKDL and CL. The first evident large gap in the existing literature is the lack of quantitative observational studies on the impact of mucocutaneous leishmaniasis on mental health. All 14 studies found examined at least one of the mental health outcomes (QoL, PM or MI). Of these studies: nine found significant associations between leishmaniasis and QoL; seven found significant associations with MI; and six with PM.

Several of the included papers appear to show an association between all forms of leishmaniasis and depression (42,43,46,49,51,52,54). Mechanisms for this are likely to be complex and interwoven as with other NTDs (62) but evidence to date suggests that scarring in CL has an increased association with social and family rejection (44) and anxiety/depression (54) and severe nodular lesions also carry an association with depression (50). The effect may be more severe in younger patients and women/girls although this needs to be investigated further. Lesions on the face have an increased association with anxiety/ depression (49,54). Evidence for a societal link was found between VL and mental health outcomes (MI, QoL, PM). Survey work assessing the perceptions and attitudes of the community in Bihar, India, found that there is social stigma attached to VL, as well as PM MI, and decrease in QoL due to financial loss (46). While this supports the hypothesis that financial burden may also lead to decreased mental well-being, our study did not find more evidence associating VL with financial problems and social drift. A more complete study would involve economic evaluations to be part of our inclusion criteria and require different search terms.

When looking at subtypes of leishmaniasis more specifically, there are several gaps in the literature concerning the effect of VL on mental health outcomes, despite it having been shown that VL did decrease the quality of life of patients directly (42,43). We found no evidence that neuroinflammation or inflammatory responses play a role in the development of mental health problems associated with leishmaniasis as this had not been specifically explored in this population. This is a surprising gap in the literature given that inflammation associated with other diseases has been shown to be associated with mental health (in a bidirectional manner), although these reported diseases and syndromes are all chronic (63–65). Neurological manifestations can occur in VL (12,66), which can both decrease quality of life and increase the likelihood of mental illness in Leishmaniasis. Only one review (non-systematic) looks at the neurological and psychological consequences of visceral leishmaniasis in humans and animals (67) but the only article that documents mental health outcomes in human VL patients was an article published by Carswell in 1953 (68) which did not meet our inclusion criteria and included no validated assessments of mental health outcomes.

The subject of stigma in VL was only examined indirectly in a single study from the 14 included studies. This showed a reluctance in the community to report the occurrence of the disease, indicating fear of the negative consequences this would have upon the patient and their families (46).

The association of VL and MI was, however, indirectly examined (46), where 43% of the community perceived that VL could cause changes in mental health, although no patient outcomes were reported. The prospective longitudinal study from Ethiopia addressed the question partly because it was related to HIV-VL co-infected patients (42,43) and not just VL infection. This study reported a higher mean score for depressive-symptoms compared to HIV-patients alone. However, it would be speculative to conclude that this was due to VL alone and it is possible that any other co-morbidity could exacerbate the symptoms of MI. Carswell reported “mental depression” and “apathy” in all of 96 (100%) patients of VL (68), but no clinical or diagnostic measures were used and these findings have therefore not been tested in more methodologically robust research using validated tools to assess MI, QoL or PM. Bennis et al., 2018 focuses on the stigmatising impact that localised cutaneous leishmaniasis can have, by dividing “stigma” into the categories of social stigma (when society rejects or excludes against people even if the stigmatized disagree with the way they are being treated) and self-stigma (the internalised mechanism within the person who is being stigmatized who faces rejection in an anticipatory manner) (41). Even though Pal et al 2017 do find a significant decrease in quality of life in the personal relationship domain (50), one cannot jump to conclusions about whether this was related to stigma, and whether PKDL patients suffered from MI. CL patients are victim to social stigma, self-stigma and suffer from PM, MI and decreased QoL (44,51,52,54). In Chahed et al., 2016, the number of body scars was weakly correlated to an experience of stigma (p=0.06, r=0.29). They also observed anticipation avoidance of stress, indicating not only social stigma but also self-stigma (44). These results should be interpreted with caution, because the sample size was small (n=41). The study showed stronger correlations between CL and loss of self-esteem and feelings of inferiority (r=0.77, p<0.05), and it was shown that at younger ages, women experienced higher levels of rejection and avoidance of stress (44). Bennis’ review included and reported on the results of qualitative studies (not included in the study design for this systematic review) showing: social isolation; social contempt; social exclusion; marriage difficulties; embarrassment; shame; sadness; disgust; shyness; and decreased marriage prospects (41). The scoping review concludes that stigma is closely linked to psychosocial morbidity (41).

Although PKDL and CL are caused by different *Leishmania* species, both have visible and disfiguring clinical manifestations. Stigma was found to play an important role in the mental health outcomes associated with CL and PKDL but the measurement tools used in the quantitative observational studies (part of inclusion criteria) could not quantify how much stigma a person or their family faces. Bennis et al., 2018 call for the development of a standardized tool to measure stigma (41). Stigma, feelings of rejection and social exclusion are a few of the social consequences of CL and PKDL that are much more easily studied in a qualitative setting, e.g. in focus groups or individually using non-structured open-ended questionnaires as was the case with (10,11,69). Overall, the clinical manifestations for PKDL and CL both led to decreased body satisfaction as well as misconceptions within society about potential disease spread. These findings complement those of Bailey et al, in their recent systematic review of the psychosocial impact of CL. They used the data from qualitative and quantitative studies to demonstrate a high burden of co-morbid depression in both active and inactive forms of the disease (70).

The only studies including a co-morbidity were part of the same study assessed at baseline (cross-sectional) (42) (prospective cohort) (43) where the impact of VL as a comorbidity of HIV was assessed pre- and post-treatment with anti-parasitic treatment for leishmaniasis. The baseline study shows that HIV-VL patients showed significantly worse quality of life scores in all domains compared to the HIV-alone patients (p=0.001)(42). Regarding psychosocial morbidity and mental illness, the Kessler Psychological Distress Scale correlated significantly with psychological health, social relation, and environmental domains of the WHOQoL had correlation coefficients of −0.335, −0.295, and −0.350 with the Kessler scale (p=0.001), further showing that a decrease in quality of life correlates with an increased mean psychological distress score. The mean (SD) depressive symptoms scale score was higher in HIV-VL patients compared to HIV patients, 2.67 (±0.7) vs 1.161 (±0.5) respectively (42). After therapy, at the 6-month follow-up stage (43), HIV-only patients showed no significantly different QoL scores compared to HIV-VL patients, showing that QoL of life scores improve after the disease is treated. Usually patients return to their normal physical appearance, as opposed to the other forms of leishmaniasis. This is starkly different in comparison to people who have CL and PKDL lesions. When the cutaneous lesions are healed, these become visible, disfiguring scars on the patient. Studies have shown that the suffering is long-term with the consequences of the disease for life (50,51,54).

We used the results of the review to develop a conceptual framework (Figure 1) outlining the relationships visually between the forms of leishmaniasis, mental illness, psychosocial morbidity and quality of life. This shows the strength of evidence or the absence of evidence where there are hypothesised links. This takes the form of a model that may help to direct future research and public health policy.

**Fig 1.**
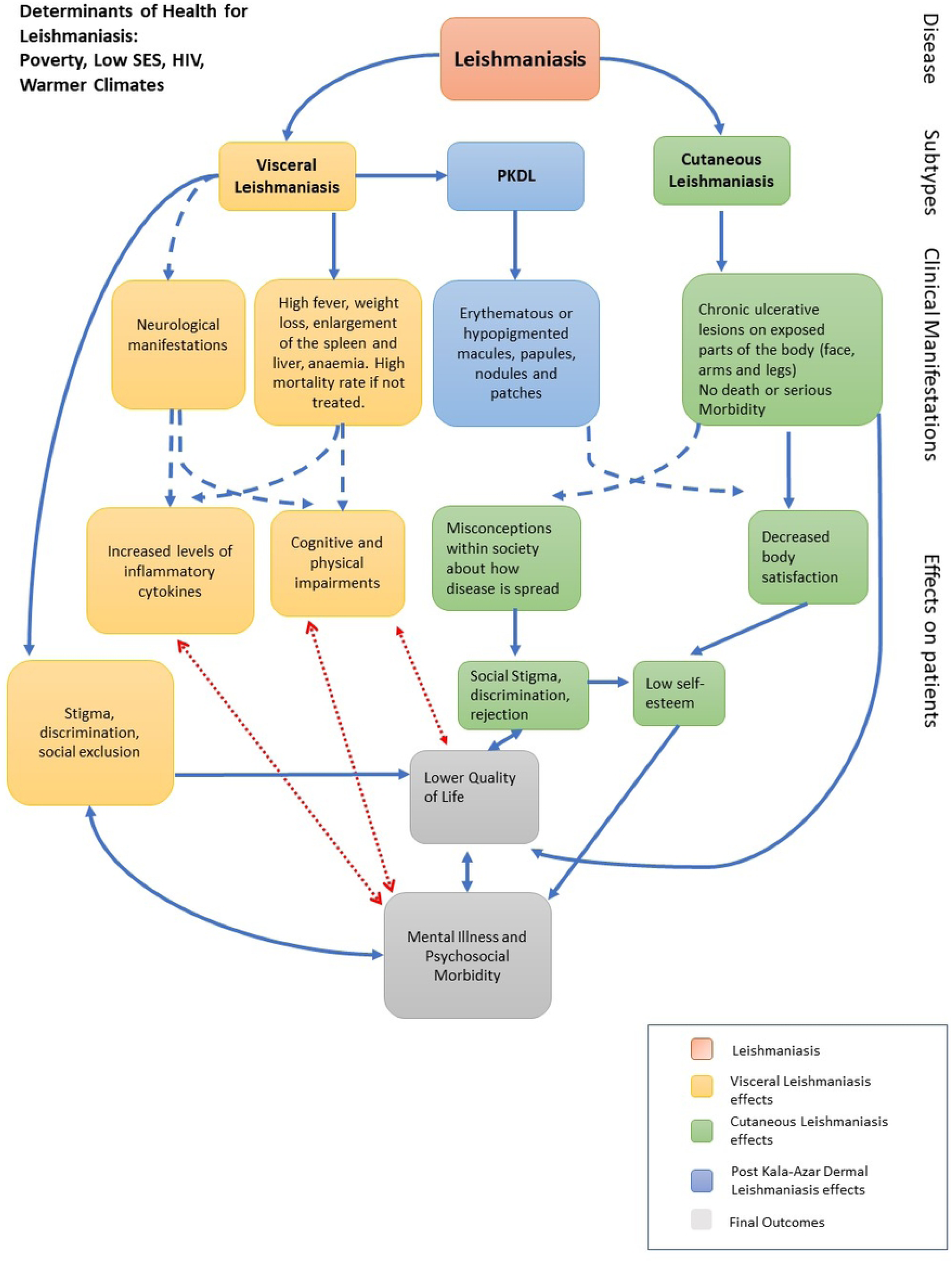
Conceptual Framework. Theoretical model linking leishmaniasis to decreased quality of life, mental illness and psychosocial morbidity. Continuous blue arrows show links that were confirmed in the included studies, dotted blue arrows show links that have been established in the literature but were not mentioned in our included studies, and red arrows show theoretical links that were hypothesized for this systematic review but not confirmed in the literature.

### Limitations

The variability in outcome measures and study designs contributed to large inter-study heterogeneity. Therefore, a narrative description of the results was the only option. Even if the outcome measures had been more consistent or if more papers had been identified, the lack of consistent good quality studies, would have hindered the conduct and interpretation of any meta-analysis. This highlights the need for better literature in this area. The searches and inclusion criteria for this review did not include qualitative studies which could also have revealed useful information regarding the studied outcomes.

### Future research

Further research is recommended with larger sample sizes. Research should include both mixed-methods and qualitative methods, as it is only through combining quantitative and qualitative measures of mental health outcomes that we will begin to understand the dimensions of the problem at hand.

There has been no research to follow-up on Carswell’s observation in 1953 (68), that most patients with VL showed marked signs of depression. Thus, a more sophisticated, and modern large cross-sectional study with VL patients to study if there is associated mental illness in these patients would be an important aspect to investigate. This research needs to be designed in culturally appropriate ways given the variety of low- and middle-income country settings that the disease is to be found.

### Clinical and Policy Implications

Governments of endemic countries could invest more in research to find out exactly what is the best way to intervene with the CL and PKDL mental health crisis at hand.

There is evidence that stigma is a serious problem associated with leishmaniasis and that for this reason policies addressing informational gaps and misinformation may be productive. Women and children were found to be impaired significantly in these studies. Women’s health is of paramount importance in public health. There is a need for effective promotion of good mental health for women and children. Hence good quality evidence-based research and practice would add considerably to this field.

In the LMIC settings, where VL is endemic, there is very little investment in mental health. If VL has a causative link with depression it may be going undiagnosed. If so, this could (after treatment) have a detrimental impact on the patients’ quality of life, and wider societal and economic impacts.

## CONCLUSION

This wide exploratory systematic review has shown that there are substantial gaps in our knowledge and in the research literature and that there is a lack of methodological quality in many of the existing studies. This systematic review shows preliminary evidence that leishmaniasis impacts upon individuals and families affecting their social status, causing stigma, with effects on quality of life and raising the risk of mental health problems. This work allows us to build a preliminary model that we present here to scaffold future attempts to better understand the effects of leishmaniasis on MI, PM and QoL.

## ACKNOWLEDGEMENTS

No funding was used to conduct this systematic review by MP, VM, BW or RC.

PK is supported by a Wellcome Trust Senior Investigator Award (WT1063203; https://wellcome.ac.uk) and MRC Global Challenges Research Fund Foundation Award (MR/P024661/1; https://www.ukri.org/research/global-challenges-research-fund/).

## Captions for Supplementary Tables

S1 PRISMA Checklist

S2 PRISMA Flow Diagram.

After deduplication of identified records, 12417 titles of records were screened. Out of 362 Abstracts that conformed to the inclusion criteria, 45 full articles were assessed for eligibility. 14 final articles were selected for analysis.

S3 Search Strategies

S4 Table 1 - Newcastle Ottawa Scale

S5 Table - ROBIS Phase 1 Assessing Relevance

S6 Table - ROBIS Phase 2 Identifying concerns about bias in the review process

S7 Table - ROBIS Phase 3: Judging Risk of Bias

S8 Table - ROBIS Phase 4: Risk of Bias in the Review

